# A novel multi-network approach reveals tissue-specific cellular modulators of fibrosis in systemic sclerosis, pulmonary fibrosis and pulmonary arterial hypertension

**DOI:** 10.1101/038950

**Authors:** Jaclyn N. Taroni, Casey S. Greene, Viktor Martyanov, Tammara A. Wood, Romy B. Christmann, Harrison W. Farber, Robert A. Lafyatis, Christopher P. Denton, Monique E. Hinchcliff, Patricia A. Pioli, J. Matthew Mahoney, Michael L. Whitfield

## Abstract

We have used integrative genomics to determine if a common molecular mechanism underlies different clinical manifestations in systemic sclerosis (SSc), and the related conditions pulmonary fibrosis (PF) and pulmonary arterial hypertension (PAH). We identified a common pathogenic gene expression signature - an immune-fibrotic axis-indicative of pro-fibrotic macrophages in multiple affected tissues (skin, lung, esophagus and PBMCs) of SSc, PF, and PAH. We used this disease-associated signature to query tissue-specific functional genomic networks. This allowed us to identify common and tissue-specific pathology of SSc and related conditions. We rigorously contrasted the lung-and skin-specific gene-gene interaction networks to identify a distinct lung resident macrophage signature associated with lipid stimulation and alternative activation. In keeping with our network results, we find distinct macrophages alternative activation transcriptional programs in SSc-PF lung and in the skin of patients with an ‘inflammatory’ SSc gene expression signature. Our results suggest that the innate immune system is central to SSc disease processes, but that subtle distinctions exist between tissues. Our approach provides a framework for examining molecular signatures of disease in fibrosis and autoimmune diseases and for leveraging publicly available data to understand common and tissue-specific disease processes in complex human diseases.

**Author Summary:** Human disease in part arises from aberrant interplay between tissues and from the interactions of gene products in tissue-specific microenvironments. Recent efforts have utilized ‘big data’ to build functional maps that model these interactions. We used these tools to study systemic sclerosis (SSc), a rare and clinically complex disease characterized by multi-organ involvement, high mortality, pulmonary fibrosis, and pulmonary arterial hypertension, and related fibrotic conditions. We developed a novel procedure to assess which processes are affected across multiple fibrotic organs and tissues. We found that patients with severe disease share molecular patterns that are indicative of dysregulated, immune and fibrotic processes. Placing these patterns into the context of functional maps allowed us to study severe disease manifestations that occur in a subset of patients. This study not only offers the potential to identify shared pathology in SSc and fibrosis, but a ‘road map’ for the use of tissue-specific networks to describe complex human diseases.

## Introduction

Integrative genomics has yielded powerful tissue-specific functional networks that model the interaction of genes in these specialized ‘microenvironments’ (1). These tools hold promise for understanding how genes may contribute to human diseases (2) that arise, in part, out of an aberrant interplay of cell types and tissues. Network biology has played a crucial role in our understanding of complex human diseases such as cancer (3,4), and more recently, in disorders where the interactions among multiple tissues are dysregulated (5).

Analytical approaches that leverage biological ‘big data’ can be especially fruitful in rare and heterogeneous diseases (6), in which the risk of mortality is significant and no approved treatments exist. We performed an integrative, multi-tissue analysis for systemic sclerosis (SSc; scleroderma), a disease for which all of these tenets are true, and included samples from patients with pulmonary fibrosis (PF) and pulmonary arterial hypertension (PAH). SSc is characterized by abnormal vasculature, adaptive immune dysfunction (autoantibody production), and extracellular matrix (ECM) deposition in skin and internal organs. The etiology of SSc is unknown, but it has complex genetic risk (7) and postulated triggers include immune activation by cancer (8), infection (9), or dysbiosis (10). SSc is clinically heterogeneous with some patients experiencing rapidly progressive skin and internal organ disease, while others have stable disease that is largely limited to skin. Understanding the drivers of disease in multiple affected organ systems is critical to understand the pathogenesis of SSc and other complications, such as PF and PAH, that co-occur in these patients.

These ‘big data’ approaches integrate individual experiments measuring hundreds of disease states and biological perturbations. Integration of these data holds promise for understanding how genes contribute to organ specific manifestations of human diseases (2). We previously developed mutual information consensus clustering (MICC) to identify gene expression that is conserved across multiple, disparate datasets (11). Here we expanded MICC to perform an integrative, multi-tissue analysis of SSc and related fibrotic conditions. Following MICC, we used the Genome-scale Integrated Analysis of gene Networks in Tissues (GIANT) tissue-specific functional genomic networks (1) to identify gene-gene interactions among those expressed consistently across affected tissues. These GIANT networks are a detailed, genome-scale representation of the functional interactions between genes in different microenvironments. We included gene expression datasets from ten different cohorts representing four different affected tissues from patients with SSc. We identified a pathogenic signature – a common ‘immune-fibrotic axis’ – that is present in all tissues analyzed and is increased in the most severe disease complications, including PF and PAH.

The immune-fibrotic axis implicates alternatively activated M∅s as central drivers of fibrosis in all solid organs studied. M∅s are highly plastic cells implicated in a wide range of pathologic processes (12-14). Using tissue-specific functional networks (1), we analyzed the nature of the immune-fibrotic axis to understand the gene-gene interactions that underlie fibrosis across organ systems. Using differential network analysis, we were able to identify skin-and lung-specific gene-gene interactions relevant to M∅ plasticity and SSc pathophysiology. We now propose a model that implicates alternatively activated M∅s as part of the immune-fibrotic axis that may drive fibrosis in multiple tissues.

## Results

We performed an integrative analysis of ten independent gene expression datasets containing samples from patients with systemic sclerosis and associated co-morbidities (Table 1). A total of 573 samples from 321 subjects recruited at seven independent centers were analyzed. These data represent samples from four different affected tissues derived from seven different clinic centers in the US and Europe. Data include SSc and control skin from a University of California, San Francisco cohort (15), a Boston University cohort (16) and a Northwestern University cohort (17). Many patients in the skin cohorts provided lesional (forearm) and non-lesional (back) skin biopsies; a subset of patients in the Northwestern skin cohort provided biopsies longitudinally over time as part of a clinical trial for mycophenolate mofetil (MMF). Peripheral blood mononuclear cell (PBMC) samples from patients with and without SSc-PAH, patients with idiopathic PAH (IPAH) and healthy controls were included from a Boston University cohort (18) and a University of Colorado PAH cohort (19). Lung data contained a cohort of late or end-stage patients that underwent lung transplant at the University of Pittsburgh (20) and a second cohort from open lung biopsies from early SSc-associated PF obtained in Brazil (21). The lung biopsies included patients with SSc-associated PF, idiopathic PF (IPF), SSc-associated PAH, and idiopathic PAH (IPAH). Data on previously unpublished samples were also included in these analyses. These are two datasets of skin biopsies from patients with limited cutaneous SSc (LSSc) recruited from University College London (UCL) / Royal Free Hospital and Boston University Medical Center. Only data that were judged to be high quality were included in the analyses. To our knowledge, there was no overlap between the patient cohorts beyond 5 patients recruited at Northwestern that provided both skin and esophageal biopsies. We summarize all patient cohorts in S1 Table.

**Table 1.** **Datasets included in this study**.

| Dataset label | Tissue | Phenotypes of interest | Citation(s) | GEO Accession |
| --- | --- | --- | --- | --- |
| Milano | Diffuse skin | Inflammatory subset, Proliferative subset | Milano et al. (15) | GSE9285 |
| Pendergrass | Diffuse skin | Inflammatory subset, Proliferative subset | Pendergrass et al. (16) | GSE32413 |
| Hinchcliff | Diffuse skin | Inflammatory subset, Proliferative subset | Hinchcliff et al. (17)<br>Mahoney et al. (11) | GSE45485, GSE59785 |
| LSSc | Limited skin | N/A | Present study | GSE76806 |
| UCL | Limited skin | N/A | Present study | GSE76807 |
| Christmann | Lung | SSc-PF | Christmann et al. (21) | GSE76808 |
| Bostwick | Lung | SSc-PF, IPF, IPAH, SSc-PAH | Hsu et al. (20) | GSE48149 |
| Esophagus | Esophagus | Inflammatory subset, Proliferative subset, SSc-PAH | Taroni et al. (75) | GSE68698 |
| PBMC | PBMC | SSc-PAH | Pendergrass et al. (18) | GSE19617 |
| Risbano | PBMC | IPAH, SSc-PAH | Risbano et al. (19) | GSE22356 |
| Abbreviations: PBMC – Peripheral blood mononuclear cells, PAH – Pulmonary arterial hypertension, PF – Pulmonary fibrosis |  |  |  |  |

The primary goal of this study was to identify the fundamental processes that occur across end-starget and peripheral tissues of patients with SSc and related fibrotic conditions. Secondly, we aimed to identify the presence or absence of common gene expression patterns that underlie the molecular intrinsic subsets of SSc (15) in different organs. Analysis of multiple tissue biopsies from patients with skin fibrosis, esophageal dysfunction, PF and PAH, allowed us to determine in an unbiased analysis whether these tissues were perturbed in a similar manner on a genomic scale.

We applied MICC (11) to identify conserved, differentially co-expressed genes across all tissues in our SSc compendium. MICC is a ‘consensus clustering’ procedure,meaning that it identifies the *shared co-clustering of genes* present in multiple datasets. MICC identifies genes that are consistently coexpressed in multiple tissues. Procedurally, MICC clusters gene expression data into coexpression modules using weighted gene correlation network analysis (WGCNA) (Fig 1). Because this clustering is purely data-driven, coexpression modules derived from different datasets necessarily differ from each other. MICC integrates these coexpression modules across datasets by identifying significant overlaps between modules from different datasets and forming a ‘ module overlap network’. MICC then parses the module overlap network to find sets of modules (communities) that are strongly conserved across many datasets (see Methods). These strongly overlapping modules correspond to molecular processes that are conserved across multiple datasets.

**Fig. 1.**
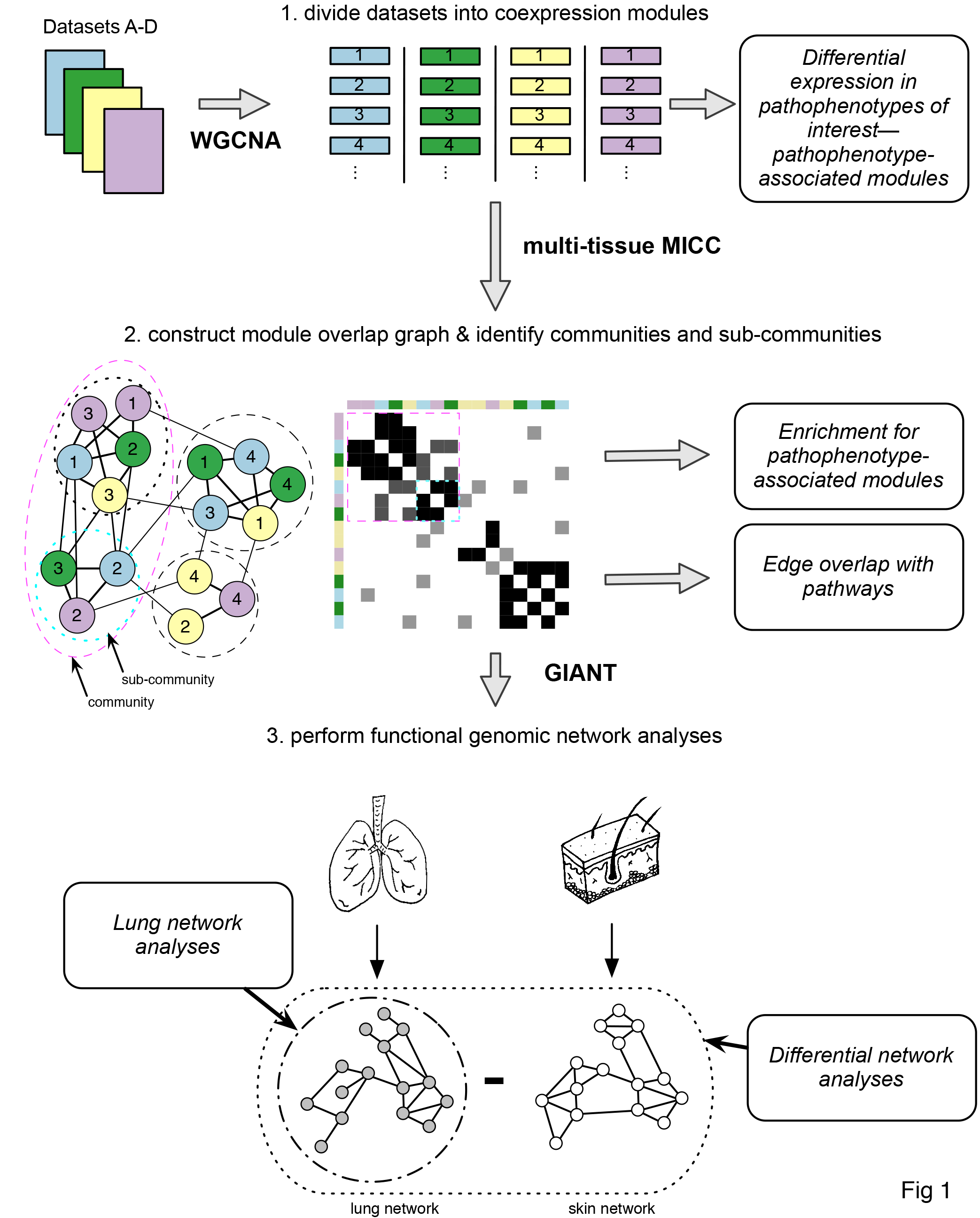
**Schematic overview of analysis pipeline**. Four datasets are shown for simplicity. Each gene expression dataset was partitioned using WGCNA independently to obtain coexpression modules. Module eigengenes were tested for their differential expression in pathophenotypes of interest. Modules were compared across datasets using MICC to form the ‘ module overlap graph’ and community detection algorithms were used to identify communities and subcommunities in the graph. These communities correspond to molecular processes that are conserved across datasets. Each community was examined for enrichment of pathophenotype-associated modules and edge overlap with canonical biological pathways. Gene sets derived from these communities were used to query GIANT functional genomic networks. The resulting networks allow for tissue-specific interrogations of the gene sets. Differential network analysis was performed to compare the lung and skin networks.

All datasets were partitioned into coexpression modules using WGCNA, resulting in 549 modules (Table 2). We constructed the 10-partite module overlap network (fig 2) and identified eight communities in the network using modularity-based community detection methods. Because the community structure of the module overlap network was hierarchical, we used a hierarchical labeling scheme, where numerals denote large communities and letters denote smaller sub-communities (Fig 2A). For each community, we used set theoretic formulae to derive a final gene set (‘consensus genes’) associated with the modules in that community (see Methods and S2 Table; consensus gene sets ranged from 64–9597 genes in size). The majority of the consensus gene sets pertain to biological processes that are not disease-specific. These include processes such as telomere organization (1A) and macromolecule localization (3A). *Disease-specific* consensus genes were identified by first determining which communities contained modules associated with pathophenotypes under study and then deriving consensus gene sets from those combined communities (see below).

**Table 2.** **Number of arrays and WGCNA coexpression modules in each of the datasets included in this study**.

| Datasets | Number of Arrays | Number of Coexpression Modules |
| --- | --- | --- |
| Milano | 75 | 39 |
| Pendergrass | 89 | 38 |
| Hinchcliff | 165 | 62 |
| LSSc | 24 | 39 |
| UCL | 15 | 98 |
| Christmann | 18 | 56 |
| Bostwick | 62 | 54 |
| Esophagus | 33 | 71 |
| PBMC | 54 | 38 |
| Risbano | 38 | 54 |

**Fig 2.**
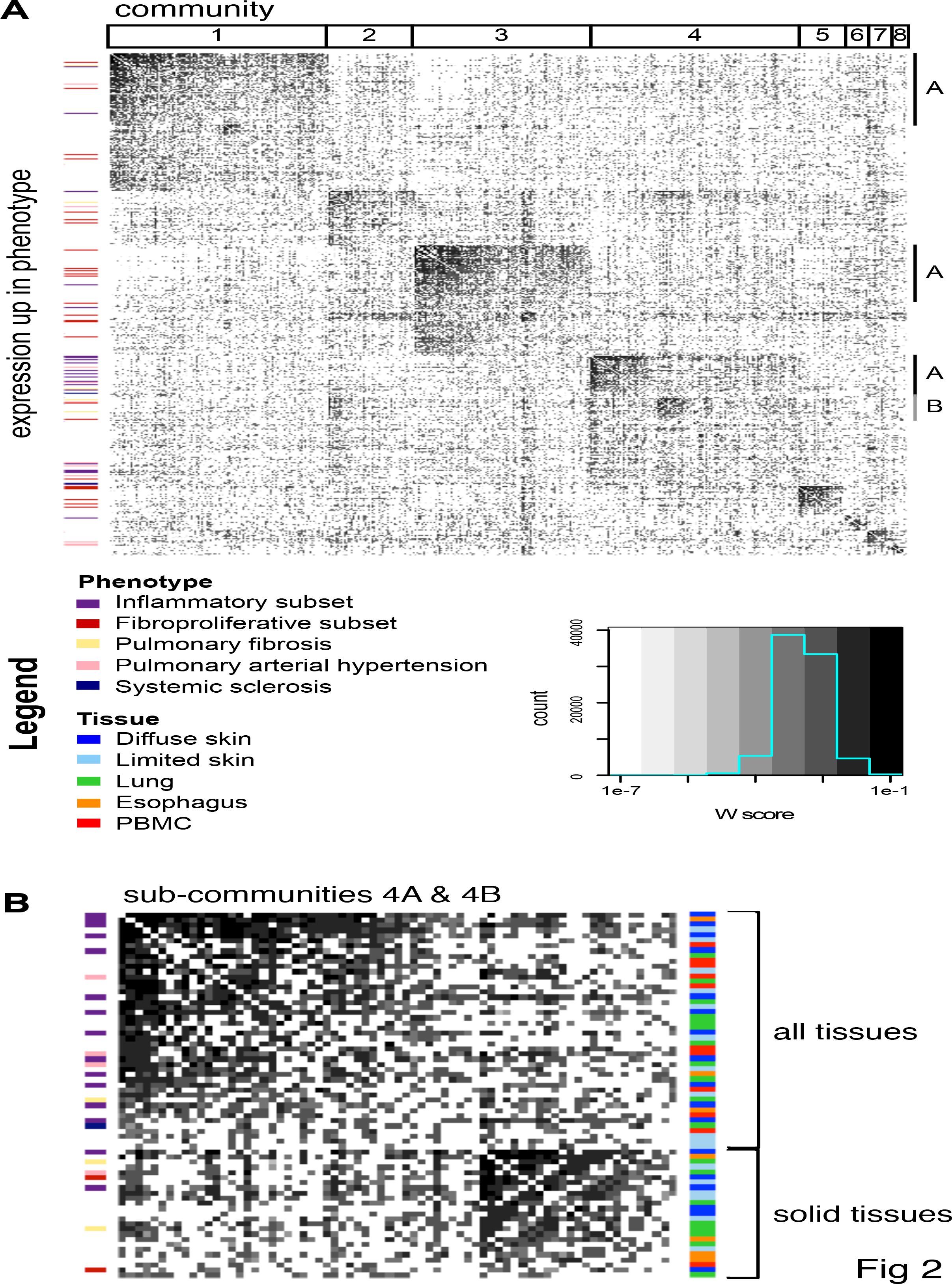
**The multi-tissue module overlap graph demonstrates that severe pathophenotypes have similar underlying expression patterns**. (A) The full adjacency matrix of the module overlap graph sorted to reveal hierarchical community structure. A darker cell color is indicative of a higher W score or larger edge weight. Communities (numbered) and subcommunities (lettered) are indicated by the annotation tracks above and on the right side of the matrix, respectively. Coexpression modules with expression that is increased in a phenotype of interest are marked by the annotation bar on the left side of the matrix. If a module was up in SSc as well as another pathophenotype of interest, the other pathophenotype color is displayed. (B) The adjacency matrix of sub-communities 4A and 4B indicates that these clusters contain modules that are up in all pathophenotypes of interest and show that there are many edges between the two sub-communities. Subcommunity 4A contains modules from all tissues whereas 4B contains mostly solid tissue modules as indicated by the tissue annotation track to the left of the matrix.

### Severe pathophenotypes share a common immune-fibrotic axis

The module overlap network is agnostic to the clinical phenotypes corresponding to each biopsy. To associate communities in the module overlap network with SSc and fibrotic pathophenotypes, we tested each of the 549 modules for differential expression in relevant pathophenotypes (see Methods). For example, every lung module in the PAH cohorts was tested for differential expression in PAH. Clusters 4A and 4B in the module overlap network contain modules with increased expression in all pathophenotypes of interest: the inflammatory and proliferative subsets of SSc, PAH, and PF (Fig 2B). Thus, the modules in these communities correspond to a common, broad disease signal that is present in every pathophenotype under study. As with our prior studies, we did not find a strong association with autoantibody subtype and the co-expression modules identified here.

Edges in the module overlap graph represent overlap between coexpression modules in different datasets, so we identified the intersection of genes between adjacent modules. We then asked if these ‘edge gene sets’ were similar to known biological processes by computing the Jaccard similarity between edges and canonical pathways from the Molecular Signatures Database (MSigDB; see Methods) (22). Edges in 4A encode immune processes such as antigen processing and presentation and cytotoxic T cell and helper T cell pathways (Table 3). This cluster also contains modules from all tissues, including PBMCs (fig 2B). Altered immunophenotypes have been reported in SSc-PAH and SSc-PF (18-21). Here, we find that the immune processes with increased expression in these severe pathophenotypes have substantial overlap with each other, as well as with the inflammatory subsets in esophagus and skin (Fig 2B and S1). Notably, 4A is composed of modules with increased expression in PAH in PBMCs and lung, and a module upregulated in end-stage PF (S1 Fig). This demonstrates a commonality of molecular pathways between the inflammatory component of SSc and the most severe end-organ complications at the expression level.

**Table 3.**
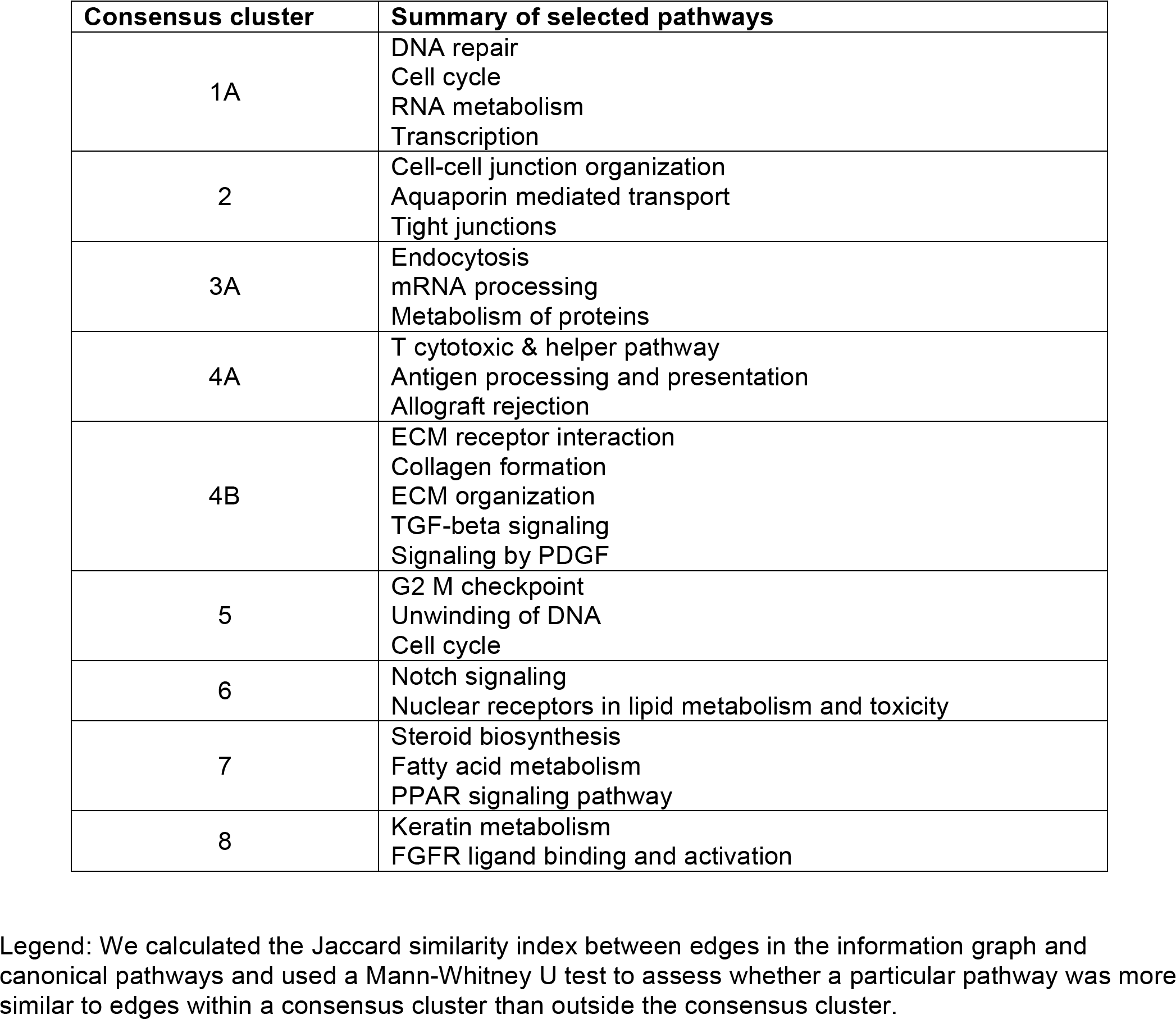
**Selected pathways that are similar to overlapping coexpression patterns in consensus clusters in the information graph**.

| Consensus cluster | Summary of selected pathways |
| --- | --- |
| 1A | DNA repair<br>Cell cycle<br>RNA metabolism<br>Transcription |
| 2 | Cell-cell junction organization<br>Aquaporin mediated transport<br>Tight junctions |
| 3A | Endocytosis<br>mRNA processing<br>Metabolism of proteins |
| 4A | T cytotoxic & helper pathway<br>Antigen processing and presentation<br>Allograft rejection |
| 4B | ECM receptor interaction<br>Collagen formation<br>ECM organization<br>TGF-beta signaling<br>Signaling by PDGF |
| 5 | G2 M checkpoint<br>Unwinding of DNA<br>Cell cycle |
| 6 | Notch signaling<br>Nuclear receptors in lipid metabolism and toxicity |
| 7 | Steroid biosynthesis<br>Fatty acid metabolism<br>PPAR signaling pathway |
| 8 | Keratin metabolism<br>FGFR ligand binding and activation |

Legend: We calculated the Jaccard similarity index between edges in the information graph and canonical pathways and used a Mann-Whitney U test to assess whether a particular pathway was more similar to edges within a consensus cluster than outside the consensus cluster.

Edges in 4B encode pro-fibrotic processes including ECM receptor interaction, collagen formation, and TGF-β signaling (Table 3). Cluster 4B consists of skin inflammatory and fibroproliferative subset-associated modules as well as lung PAH-, late PF-and early PF-associated modules (Fig 2B and S1). These results validate and expand what we have found in our prior meta-analysis of skin data alone (11): the immune-fibrotic axis observed in the SSc intrinsic subsets are *connected* to and, furthermore, *are found in all* other tissues and SSc-associated pathophenotypes.

To understand how the immune-fibrotic axis and these phenotypes are functionally related, we identified the consensus genes in the combined 4A and 4B clusters (see Methods; 2079 unique genes; S4 Table). Using a conservative measure, these consensus genes are enriched for genes with increased expression in all disease manifestations (Significance Analysis of Microarrays or SAM (23), FDR <5%) (PF in both lung datasets p < 2.2e–16; PAH lung, p = 7.88×10^−5^; PAH in both PBMC datasets, p = 3.20×10^−15^, Fisher’s exact test). This demonstrates that the tissue consensus genes are highly relevant to all disease manifestations in this study. The tissue consensus gene sets allow us to rigorously extrapolate from this conservative set a substantially broader, disease-associated signal. This extrapolation is especially important for tissue studies that are underpowered to detect a large number of significantly differentially expressed genes (see Discussion). We took the union of the tissue consensus gene sets as a set of ‘immune-fibrotic axis consensus genes’ that are informative about pathology in every tissue.

### The lung functional genomic network reveals a coupling of immune and fibrotic processes

The GIANT functional networks infer functional relationships between genes by integrating publicly available data including genome-wide human expression experiments, physical and genetic interaction data, and phenotype and disease data (1). In these networks, genes are nodes and edges are weighted by the estimated probability of a tissue-specific relationship between genes. GIANT contains networks for multiple tissues, including skin and lung. To investigate the function of the immune-fibrotic axis consensus genes in pulmonary manifestations of SSc, we extracted the subnetwork of the GIANT whole genome lung network corresponding to the immune-fibrotic axis consensus genes - the *lung network* (Fig 3 and S3). Similar to our previous analysis of SSc skin, we find interconnected functional modules related to both immune (interferon (IFN)/antigen presentation and innate immune/NF-κB/apoptotic processes) and fibrotic (response to TGF-β and ECM disassembly/wound healing) processes (fig 3A). This demonstrates that, like skin, there is functional coupling between inflammatory and pro-fibrotic pathways in lung.

**Fig 3.**
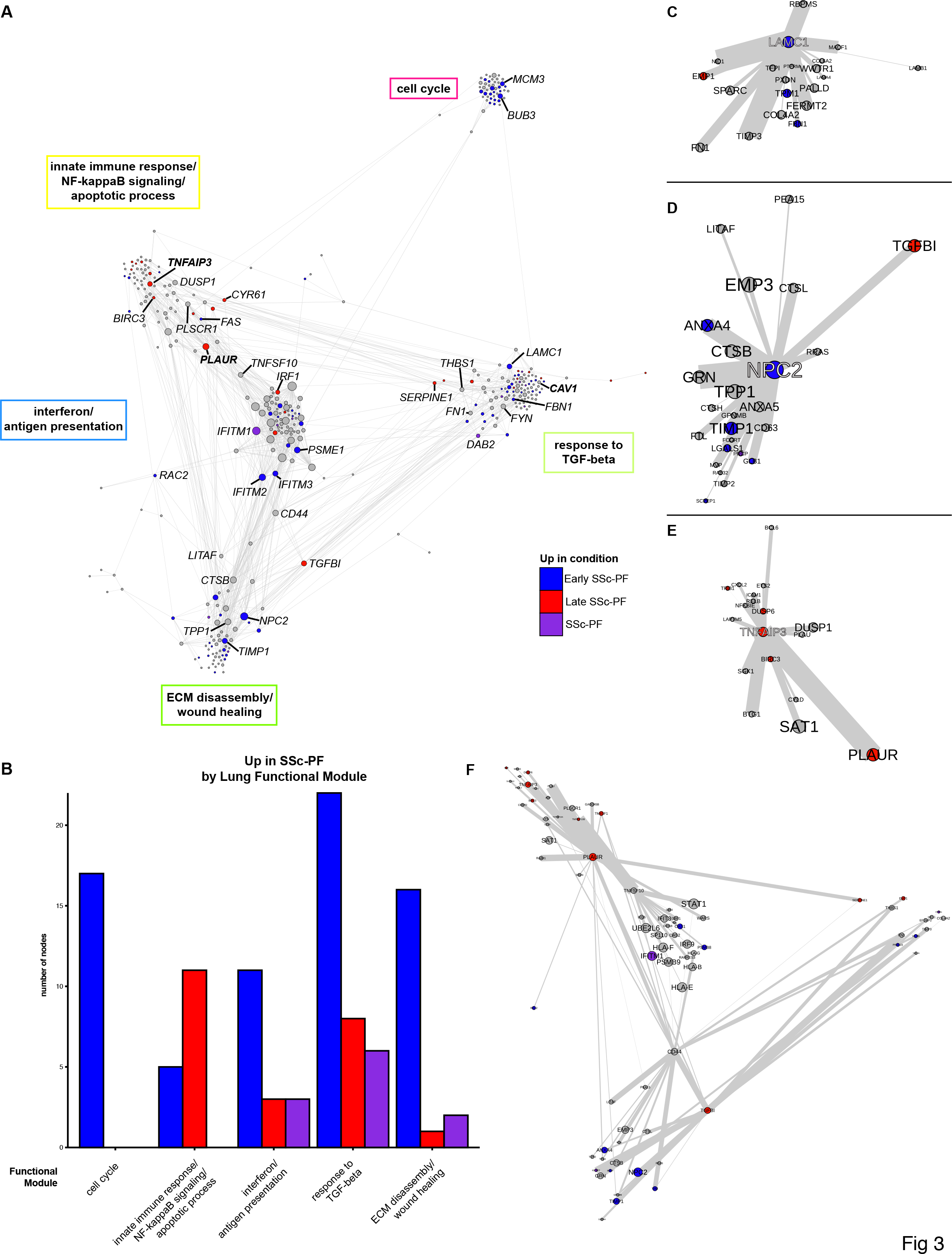
**Genes that are overexpressed in late and early SSc-PF are distributed throughout the consensus lung network**. (A) The lung network shows functional connections between inflammatory and fibrotic processes. Genes in the largest connected component were clustered into functional modules using community detection. Biological processes associated with the functional modules are in boxes next to the modules. Genes are colored by whether they are over-expressed in late SSc-PF (red), early SSc-PF (blue), both (‘SSc-PF’, purple), or neither if they are grey. Gene symbols in bold have putative SSc risk polymorphisms. Node (gene) size is determined by degree (number of functional interactions) and edge width is determined by the weight (probability of interaction between pairs of genes). The layout is determined by community membership, the strength of connections between communities, and finally the interactions between individual genes in the network. A fully labeled network is supplied as a supplemental figure intended to be viewed digitally (S3). (B) Quantification of differentially expressed genes in each of the five largest functional modules. C-E. Hubs of the consensus lung network; only the first neighbors of the hub that are in the same functional module are shown. (C) *LAMC1* is a hub of the response to TGF-beta module. (D) *NPC2* is a hub of the ECM disassembly, wound healing module. (E) *TNFAIP3* is a hub of the innate immune response, NF-κB signaling, and apoptotic processes module. (F) Bridges of the consensus lung network. First neighbors of *PLAUR, CD44, TNFSF10,* and *TGFBI* are shown.

### The lung network distinguishes early and late events in SSc lung disease

Our analysis includes two lung datasets derived from both early SSc-PF (open lung biopsies obtained for diagnostic purposes (21) and end-stage or late disease (SSc-PF patients that underwent lung transplantation (20)). In addition to the differences in disease stage between these two datasets, there is also some difference in the histological patterns of fibrosis in these cohorts. In the Bostwick lung dataset (20), all patients with SSc-PF had usual interstitial pneumonia (UIP). This study used lung tissues from patients who underwent lung transplantation (late disease). The Christmann lung dataset (21) contains 5 patients with non-specific interstitial pneumonia (NSIP) and 2 patients with centrilobular fibrosis (CLF). This study looked at early SSc-PF patients, used open lung biopsies, and specifically avoided honeycombing areas.

Although NSIP and UIP have distinct clinical outcomes, they have been shown to be nearly indistinguishable at the gene expression level (24). Furthermore, these datasets have overlapping coexpression patterns as demonstrated by their shared community membership in the module overlap network. Comparison of different datasets allows us to determine how genes with increased expression at these different stages and histological subtypes of lung disease are distributed throughout the lung network and to suggest an order of molecular events in SSc-PF progression. Genes overexpressed in SSc-PF (SAM, PF vs. Normal comparison, FDR < 5%) are distributed throughout the lung network and therefore are predicted to participate in all of the molecular processes identified in the network. Quantification of the distribution of SSc-PF differentially expressed genes throughout the consensus lung network (fig 3B) demonstrates that molecular processes can be associated either with a disease stage or transition between stages. The cell cycle module contains only early SSc-PF genes, the innate immune response/NF-κB/apoptotic processes module contains more late SSc-PF genes, and the response to TGF-β module contains genes from *both* disease stages (Fig 3A–B).

### Hub and bridge genes are highly relevant to the pathogenesis of pulmonary fibrosis

Certain genes occupy privileged positions within molecular networks and these genes often have critical biological function (25). *Module hub genes* are connected to a significant fraction of genes within a functional module, whereas *bridge genes* are genes that connect to multiple functional modules and thus ‘bridge’ them. We identified the hub and bridge genes within the lung network for their possible roles in PF pathogenesis. We highlight the hubs and bridges of the lung network in Fig 3C–E and Fig 3F, respectively. The hubs of several of the functional modules in the consensus lung network show increased expression at different disease stages (Fig 3C–E). For instance, *LAMC1* shows increased expression in early SSc-PF and is highly connected within the response to TGF-β module (fig 3C). The gene Niemann-Pick disease, type C2 *(NPC2)* is upregulated in early disease and is connected to cathepsins L and B *(CTSL, CTSB)* and *GLB1* in the lung network (fig 3D). We tabulate information on selected genes from the lung network in Table 4.

**Table 4.**
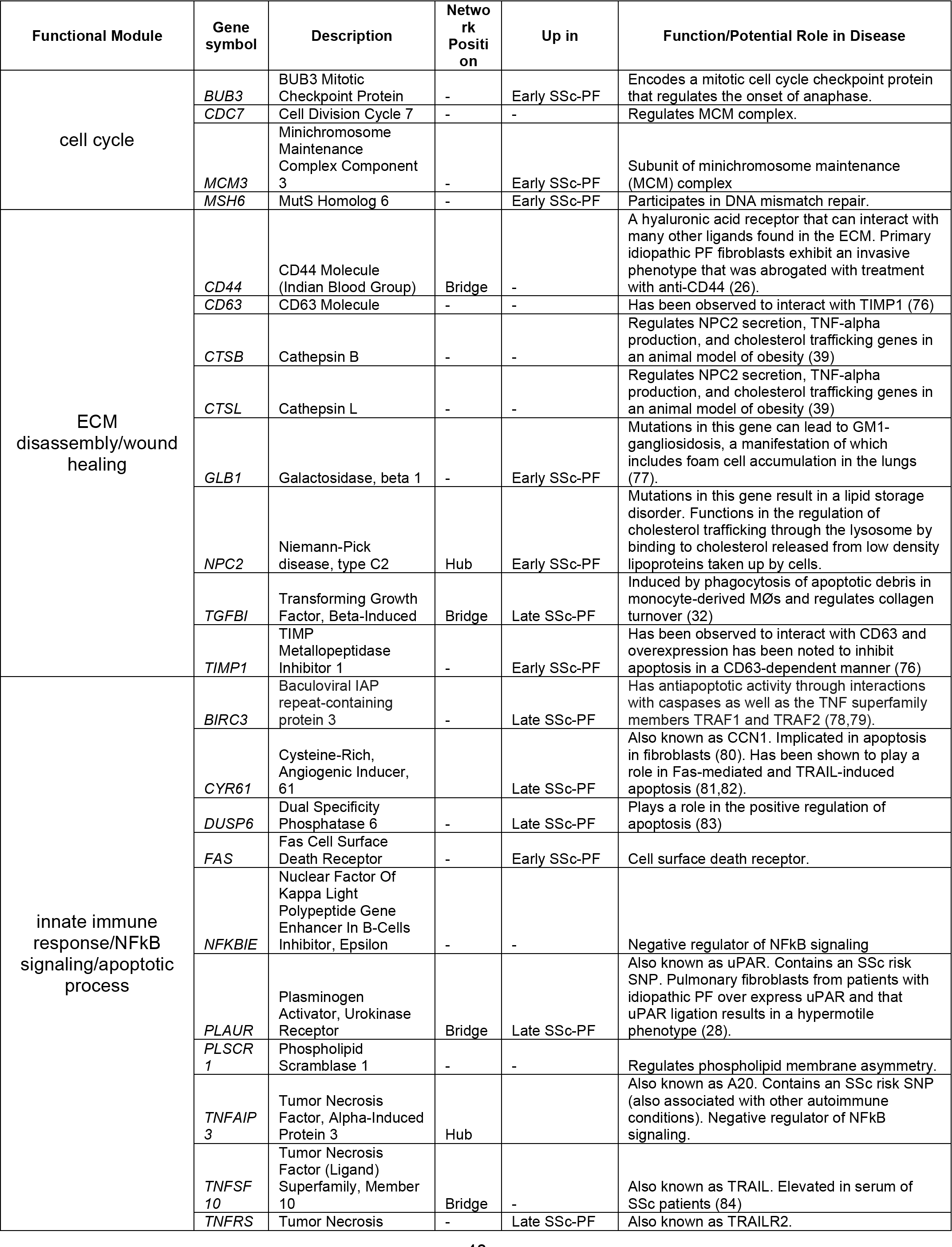
**Selected genes in the consensus lung network**.

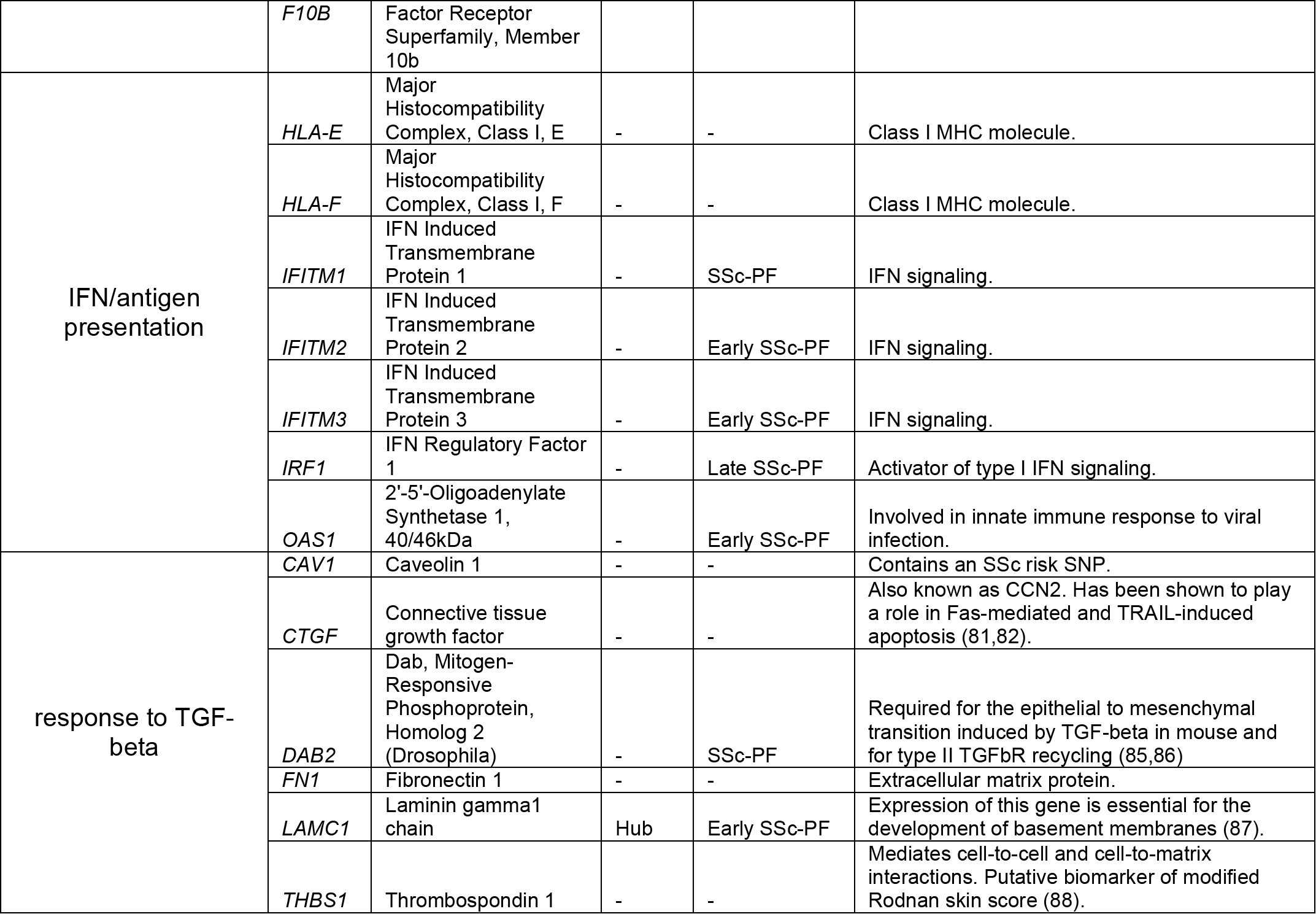

The innate immune response/NF-κB signaling/apoptotic process module contains genes that are highly expressed in late SSc-PF, including the hub genes *CYR61* and *TM4SF1* (Fig 3A–B and S3). The hub gene *TNFAIP3* (A20), which is increased in late SSc-PF (fig 3E), is a negative regulator of NF-κB signaling and inhibitor of TNF-mediated apoptosis. The innate immune response/NF-KB signaling/apoptotic process and IFN/antigen presentation modules are bridged by *TNFSF10,* also known as TRAIL (TNF-related apoptosis inducing ligand, Fig 3F). These results suggest that the balance of apoptosis is altered in late SSc-PF. The upregulation of genes with anti-apoptotic function was not reported in the original study (20), which demonstrates the strength of both the MICC method and the study of functional interactions.

*CD44* and *PLAUR* (uPAR) bridge multiple functional modules in the lung network (fig 3F) and have been implicated in IPF (26,27). Because these genes link modules important in regulating disease progression, therapeutic targeting of CD44 and μPAR may be an effective strategy in combatting SSc-PF. Indeed, anti-CD44 treatment reduces fibroblast invasion and bleomycin-induced lung fibrosis (26), and inhibition of uPAR ligation significantly reduces motility of pulmonary fibroblasts from patients with idiopathic PF (28). These results are consistent with our identification of these genes as key genes in the lung network.

### The lung microenvironment provides a distinct milieu for pro-fibrotic processes

Pulmonary fibrosis is histologically distinct from skin fibrosis and occurs in a subset of patients with SSc. We hypothesized that the lung microenvironment may have a distinct organization of immune-fibrotic axis consensus genes when compared to skin. Indeed, for interactions (edge weight > 0.5) that are present in both the lung and skin networks, there are gene pairs that are much more likely to interact in one tissue than the other (fig 4A). In other words, the skin and lung networks are ‘wired differently’. To identify *highly lung-specific* and *highly skin-specific interactions,* we performed a differential network analysis that identified gene pairs that are strongly predicted to interact in one tissue but not the other (see Methods).

**Fig 4.**
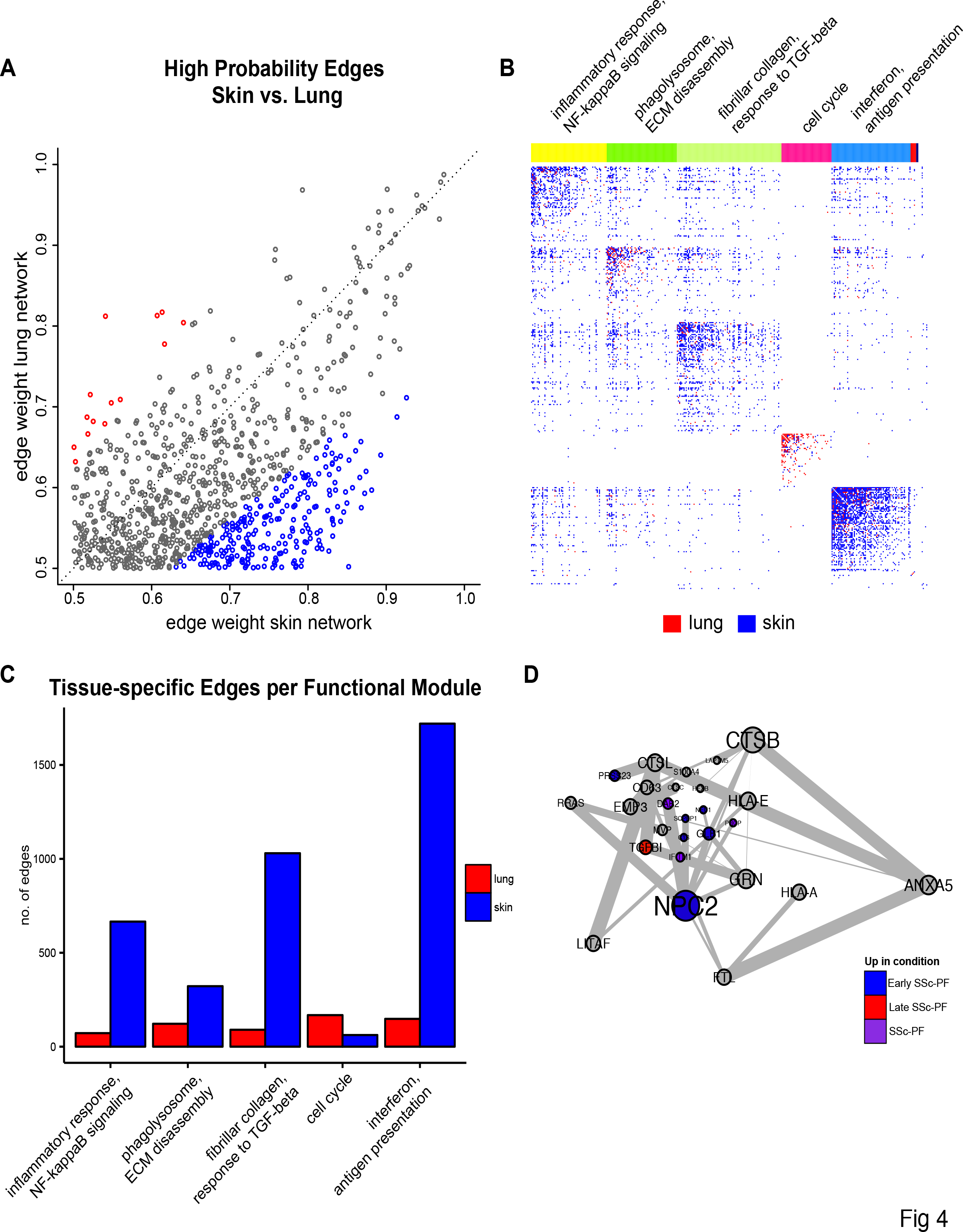
**The lung and skin network structures indicate distinct tissue microenvironments influence fibrosis**. The skin and lung networks were compared by first finding the giant component of the lung network and then collapsing to nodes only found in both the skin and lung networks (which are termed the common skin and common lung networks). (A) A scatterplot of high probability edges (> 0.5 in both networks) illustrates that pairs of genes with a higher probability of interacting in skin than lung exist and vice versa. Edges are colored red if the weight (probability) is 1.25× higher in lung or blue if it is 1.25× higher in skin. (B) The differential adjacency matrix where a cell is colored if the edge weight in a given tissue is over and above the weight in the global average and tissue comparator networks. For instance, a cell is red if the edge weight was positive following the successive subtraction of the global average weight and skin weight. Community detection was performed on the common lung network to identify functional modules; common functional modules largely recapitulate modules from the full lung network. Representative processes that modules are annotated to are above the adjacency matrix. The annotation track indicates a genes functional module membership. Nodes (genes) are ordered within their community by common lung within community degree. A fully labeled heatmap is supplied as a supplemental figure intended to be viewed digitally (S4). (C) Quantification of tissue-specific interactions in each of the 5 largest functional modules. (D) The lung-resident M∅ module found in the differential lung network (consists only of edges in red in panel B).

These highly specific interactions are displayed in Fig 4B, where a cell is red if it is lung-specific or blue if it is skin-specific (cf. S4 Fig). The number of tissue-specific edges in each functional module is quantified in Figs 4B and 4C, which illustrate that most functional modules in lung have fewer interactions than in skin, with the exception of the cell cycle module. Of particular interest is the relationship between the phagolysosome/ECM disassembly genes and response to TGF-β genes, as strong differential connectivity can be observed in this module (Figs 4B and 4C). Thus, even though ECM disassembly and TGF-β module genes are coordinately differentially expressed in both lung and skin, they are differentially connected to each other suggesting that the microenvironment strongly determines the functional consequences of upregulating these pro-fibrotic genes.

To summarize lung-specific biological processes in the immune-fibrotic axis, we clustered the lung-specific interactions (differential lung network) to identify lung-specific pathways (S5 Fig). We identified 23 clusters corresponding to biological processes such as type I IFN signaling (cluster 10), antigen processing and presentation (cluster 4), REACTOME Cell surface interactions at the vascular wall (cluster 22), and mitotic cell cycle (cluster 16, shown in Fig S5B). Taken together, this suggests that within the immune-fibrotic axis we find innate immune and cell proliferation processes that are highly lung-specific. One of the largest of these clusters (cluster 13, Fig 4D and S5C) includes *NPC2, S100A4,* and *CTSB,* which encode protein products that are highly expressed in normal lung-resident M∅s (LR-M∅s) (29,30).

*NPC2,* is a hub of the ECM disassembly/wound healing module in the full lung network (fig 3); many of the genes in cluster 13 also belong to the ECM disassembly/wound healing module in the whole network, including the cathepsins *CTSB* and *CTSL.* Alveolar M∅s are the main source of cathepsins in bleomycin-induced fibrotic lung tissue (31). Additional genes associated with development and maintenance of alternative M∅ activation include *TGFBI* (32), *NEU1* (33), *PRCP* (34), and *DAB2* (35). Genes that are specifically associated with alternative activation of lung M∅s include *PLP2* (36) and *IFITM1* (37) (Fig 4D and S5C). Based on these genes and the complete lung network in Figure 3, we identified an LR-M∅ signature. These findings are consistent with previous reports of alternative M∅ activation in SSc (21,38).

To explore this signature further, we examined some genes from this cluster along with genes identified in the Christmann, et al. study (21). Consistent with the primary publication (21), some heterogeneity in SSc-PF gene expression is observed and is likely due to tissue sampling from various lobes of the lung as well as the inclusion of patients with centrilobular fibrosis (Fig 5A, right dendrogram branch). Nevertheless, the LR-M∅ signature comprises genes that are highly correlated with canonical markers of alternatively activated M∅s that were validated by either PCR or immunohistochemistry in the original study (e.g., *CD163* and CCL18) (21).

The LR-M∅ cluster in the differential lung network also contains a number of genes implicated in lipid storage disorders, including *HEXB, GLB1,* and *NPC2.* Several other LR-M∅ cluster genes have been shown to be important for regulating cholesterol trafficking genes in an animal model of obesity, including *CTSB, CTSL,* and *NPC2* (39). It has been noted that lipid metabolism genes are upregulated in lung M∅s relative to other tissue-specific M∅s (36). Furthermore, in the bleomycin injury mouse model of pulmonary fibrosis, lipid-laden M∅s have been observed to increase expression of markers associated with alternative M∅ activation and to secrete TGF-β (40).

### Distinct M∅ gene expression programs are elevated in lung and skin

We hypothesized that early SSc-PF lung samples may have evidence of both alternatively activated and lipid-stimulated M∅s and that this may differ from what is observed in skin. The presence of alternatively activated M∅s in the inflammatory subset of skin was inferred in our single tissue analysis (11). To test this hypothesis, we used gene sets associated with classical activation of M∅s, alternative activation of M∅s, or stimulation of M∅s with a variety of activation stimuli, including free fatty acids, taken from Xue, et al. (12). To summarize the expression of each M∅ gene set (12) and compare across tissues in these data, we computed the average expression of all genes in each gene set (see Methods; see S5 Table for a mapping between Xue, et al. modules and our naming scheme). Results are displayed for control and SSc-PF lung, as well as control and SSc-inflammatory skin (fig 5B). As shown in Figure 5B, there is evidence of an increase in alternatively activated and free fatty acid stimulated gene sets in SSc-PF and SSc-inflammatory skin. These data do not show statistically significant differences in expression of gene sets associated with classical M∅ activation between controls and SSc-PF or SSc-inflammatory skin (see S6 Table for p-values of all modules tested).

**Fig 5.**
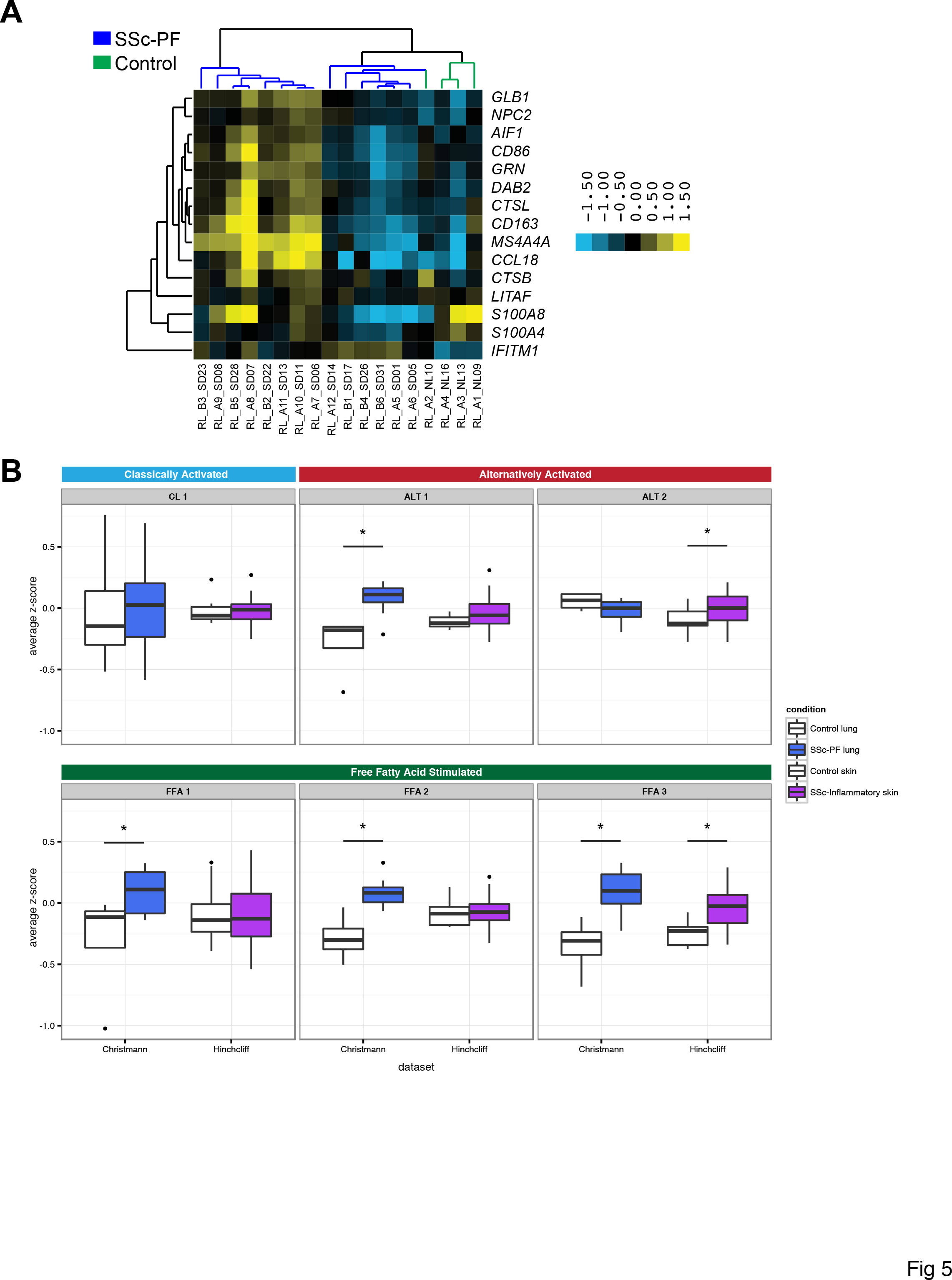
**Evidence for alternative activation of M∅s in SSc-PF lung that is distinct from**. (A) Genes identified by differential network analysis and inferred to be indicative of lung-resident M∅s are correlated with canonical markers of alternatively activated M∅s such as *CCL18* and *CD163* in the Christmann dataset. (B) Summarized expression values (mean standardized expression value) of gene sets (coexpression modules) upregulated in various M∅ states from the Christmann and Hinchcliff datasets - Module CL1: classical activation (IFN-γ); Modules ALT 1 and 2: alternative activation (IL-4, IL-13); Modules FFA 1, 2, and 3: treatment with free fatty acids. Taken from (12).

The discovery of IFN (IFN)-related genes among the consensus genes indicates that these pathways are increased in pathophenotypes of interest (e.g., SSc-PF and the skin inflammatory subset). Christmann, et al. also noted a strong IFN-related gene signature in SSc-PF samples, although the cellular compartment responsible for this signature was not described (21). Because stimulation with IFN results in classical activation of M∅s, we examined the expression of genes from CL 1, as it is most strongly associated with IFN-γ treatment (“classical activation”) in human M∅s (12). However, CL 1 genes’ expression is not different between disease and controls in either skin or lung (Wilcoxon *p* = 0.76 and 0.80, respectively; Fig 5B). This result is consistent with our inability to discern differences in classical M∅ activation markers between controls and SSc-PF and inflammatory skin and suggests that classically activated M∅s are not the source of the reported IFN signature.

Modules ALT 1 and ALT 2 are both associated with IL-4 and IL-13 treatment, which are stimuli associated with alternative activation of M∅s (12). These two gene sets are non-overlapping coexpression modules and therefore represent two “parts” of the alternatively activated M∅ transcriptional program. We performed functional enrichment analysis for ALT 1 and 2 to understand which biological processes underlie these transcriptional signatures (see Methods). Module ALT 1 is enriched for genes involved in oxidative phosphorylation (KEGG, p < 0.0001) and the citric acid cycle (REACTOME, p < 0.0001) pathways. In lung, ALT 1 expression is higher in SSc-PF than in controls (Wilcoxon *p* = 0.0046). There is no difference between healthy controls and the inflammatory subset in skin (Wilcoxon *p* = 0.41). Module ALT 2 shows an opposite trend is enriched for genes implicated in the positive regulation of response to wounding (GO BP, *p* = 0.027) and defense response (GO BP, *p* = 0.00035); this module includes alternatively activated M∅ markers such as *CD14* and *CCL26* (41,42). ALT 2 expression is increased in the inflammatory subset in skin (Wilcoxon *p* = 0.041) and trends toward decreased expression in SSc-PF lung (Wilcoxon *p* = 0.16). Together, these pathways suggest a metabolic “switch” associated with alternative activation in lung that is not found in skin (for review see (43); Fig 5B).

We also analyzed modules associated with free fatty acids (FFA) stimulation, which are relevant to the question of lipid signaling or exposure in SSc tissues (FFA 1, 2, and 3). We first performed functional enrichment analysis for these modules to gain biological insight into these transcriptional programs. FFA 1 is enriched for genes involved in the Unfolded Protein Response (REACTOME, *p* = 0.025). FFA 2 is enriched for Antigen processing-Cross presentation genes (REACTOME; p = 0. 00101). FFA 3 is enriched for genes in the ER-Phagosome Pathway (REACTOME, *p* = 0.0076). Expression of FFA 1 and 2 is significantly increased in lung (FFA 1: Wilcoxon *p* = 0.046; *p* = 0.97 in skin; FFA 2: Wilcoxon *p* = 0.0013; *p* = 0.63 in skin), whereas FFA 3 is upregulated in SSc-PF lung (Wilcoxon, *p* = 0.0013) and the SSc inflammatory subset in skin (Wilcoxon, *p* = 0.00056). These results suggest that LR-M∅s may have a distinct lipid exposure that strongly diverges from that in skin.

The differential network analysis (Fig. 4) allowed us to identify highly lung-specific interactions in the immune-fibrotic axis that implicated lipid signaling as a distinct functional process in lung. The higher expression of *multiple* free fatty acid-associated modules in lung suggests that the role of lipid signaling in M∅s may be more important in this tissue than in skin, consistent with what we would predict based on *highly lung-specific* gene-gene interactions, and based on prior biomedical literature in related conditions (36,40). Thus, a major difference between the lung and skin networks can be attributed to the presence of a distinct M∅ phenotype in lungs.

## Discussion

SSc is a systemic disease that affects multiple internal organs but, to our knowledge, no one has shown if there are distinct or common deregulated pathways between these organ systems, or their relationship to other fibrotic conditions. In recent years, gene expression data have been collected for multiple tissues. However, these data often have issues that are common to many diseases. First, SSc is rare and patients with particular disease manifestations are still rarer, so there is a limit to the amount of biopsy material available for study. Second, for practical and ethical reasons, internal organ biopsies are seldom taken from healthy subjects making comparisons difficult. Thus, lung, esophagus, and other affected internal organs are more difficult to study than blood and skin tissue. Therefore, there is a critical need to leverage our biological prior knowledge with our understanding of well-studied tissues - like blood and skin - to make plausible inferences about pathogenesis in tissues that are more difficult to study.

The clinical heterogeneity of SSc, particularly the difficulty of predicting internal organ involvement, raises an important question: are the fibrotic processes observed in multiple organs derived from a common disease process, or is each organ manifestation effectively a distinct disease? Our analyses demonstrate that there is a common gene expression signature underlying all severe organ manifestations of SSc - the immune-fibrotic axis - in solid organs. The immune-fibrotic axis underlies both SSc pulmonary manifestations of PF and PAH, and the intrinsic subsets of skin and esophagus. Moreover, coexpression modules from peripheral blood, a mixture of innate and adaptive immune cells, have significant overlap with modules associated with all pathophenotypes studied. Thus, while fibrotic processes were largely associated with solid tissues, the inflammatory component of the immune-fibrotic axis is only found in peripheral blood.

The presence of a common gene expression signature across multiple tissues suggests a common disease driver, but it does not resolve the possible tissue-specific processes that contribute to disease in the internal organs. Indeed, there are many layers of biological regulation between gene expression and whole tissue phenotypes. Resolving the relationship between molecular profiles and phenotypes is a difficult biological problem underlying most biomedical inquiry. However, these relationships have been approximated by integrating high-throughput genomic data into tissue-specific functional networks using ‘big data’ machine learning strategies (1). We addressed tissue-specificity in SSc pathology by interpreting the common expression signal - the immune-fibrotic axis - within these tissue-specific functional networks. These networks allowed us to identify critical genes that occupy important positions in molecular pathways in lung. It is clear from this work that the coupling of immune and fibrotic processes is a hallmark of SSc that occurs in SSc-PF and SSc-PAH as well as skin. However, we also find subtle, lung-specific functional differences that we attribute, in part, to the plasticity of the myeloid cell lineage.

### The plasticity of the myeloid lineage may drive tissue-specific SSc disease processes

By performing a combined analysis of SSc gene expression in multiple tissues, we are able to observe and infer, in a genome-wide manner, commonalities in the complex mixture of cell types in a tissue at the time of biopsy. Overwhelmingly, we detected a M∅ signature associated with severe disease. In the module overlap network, we find that PAH-associated modules from PBMCs (18,19) have significant overlap with SSc inflammatory subset-associated modules from skin and esophagus (fig 2). Indeed, in Pendergrass et al. *(18),* we observed that PBMCs from lcSSc patients have significant enrichment in myeloid-and M∅-related gene sets as compared to healthy controls. Christmann et al. (44) expanded on this, showing that highly expressed transcripts in lcSSc-PAH CD14^+^ monocytes were induced in IL-13-stimulated cells, i.e. that PAH monocytes are alternatively activated. We assert that this M∅ polarization is a significant part of the immune-fibrotic axis we find in these data and, therefore, is likely *a common driver* of the complex pathophysiology of SSc. In support of this, an independent study also identified M∅s and dendritic cells (DCs) as possible sources of an “inflammatory” signature in lesional SSc skin (45).

We found evidence for the contribution of LR-M∅s to SSc-PF pathobiology, consistent with the alternative activation of M∅s and TGF-β production. In our prior analysis of skin, we inferred alternatively activated M∅s as modulators of the SSc inflammatory intrinsic subset in skin (11). Our current study identifies a LR-M∅ signature within the functional relationships of immune-fibrotic axis consensus genes in lung (Fig 4D and 5A). We posit that the differences in fibrotic responses of skin and lung tissue are due, in large part, to innate differences between tissue-resident M∅s that have been observed (46,47), as well as the interactions between infiltrating monocytes and tissue-resident cell types (e.g., alveolar epithelial cells vs. keratinocytes). Because M∅ phenotype and function are plastic and readily modulated by the local tissue microenvironment, it is likely that differential activation of M∅s in these tissues is the result of exposure to distinct cytokine milieu. Indeed, we show that distinct alternative activation gene expression programs have increased expression in SSc-PF lung and inflammatory SSc skin (Fig. 5). In particular, there were multiple lipid-related signatures elevated in SSc-PF lung alone.

We cannot rule out that the M∅ changes we observe are a secondary response to the affected organ pathology. Regardless, therapies that target M∅ effectors such as IL6R have shown promise in clinical trials (48) and M∅ chemoattractants have been shown to be important in animal models of SSc inflammatory disease (49), suggesting that M∅s play a central role in SSc pathogenesis. We also cannot rule out that DCs contribute to our results, as plasmacytoid DCs are observed to be important in the Stiff Skin Syndrome mouse model (50). However, some skin-resident DCs have been shown to be transcriptionally similar to peripheral blood monocytes in humans (51). We speculate that the circulation of peripheral myeloid cells contributes to the multi-organ nature of SSc. Future studies may use *in silico* and cell sorting techniques to deconvolve SSc expression data to identify changes in cell proportion and transcriptome throughout disease course and to finely phenotype myeloid cells from SSc patient tissue samples.

### An overview of SSC-PF disease processes

The study of two different lung datasets that sampled early-and late-stage SSc-PF allows us to describe differences between the disease processes found in these two datasets. The two datasets each contained patients with different types of interstitial pneumonia (see Methods), which may limit interpretation of these results. However, as stated in the results, we and others (24) find evidence of highly similar gene expression patterns between UIP and NSIP. We do not have treatment information for patients in these studies and acknowledge that late-stage patients are more likely to be treated with immunosuppressive therapy. With these caveats in mind, we can nevertheless draw non-intuitive conclusions through the combination of our data-driven approach and mechanistic insight from disparate literature. We provide an overview of disease processes in Fig 6.

**Fig 6.**
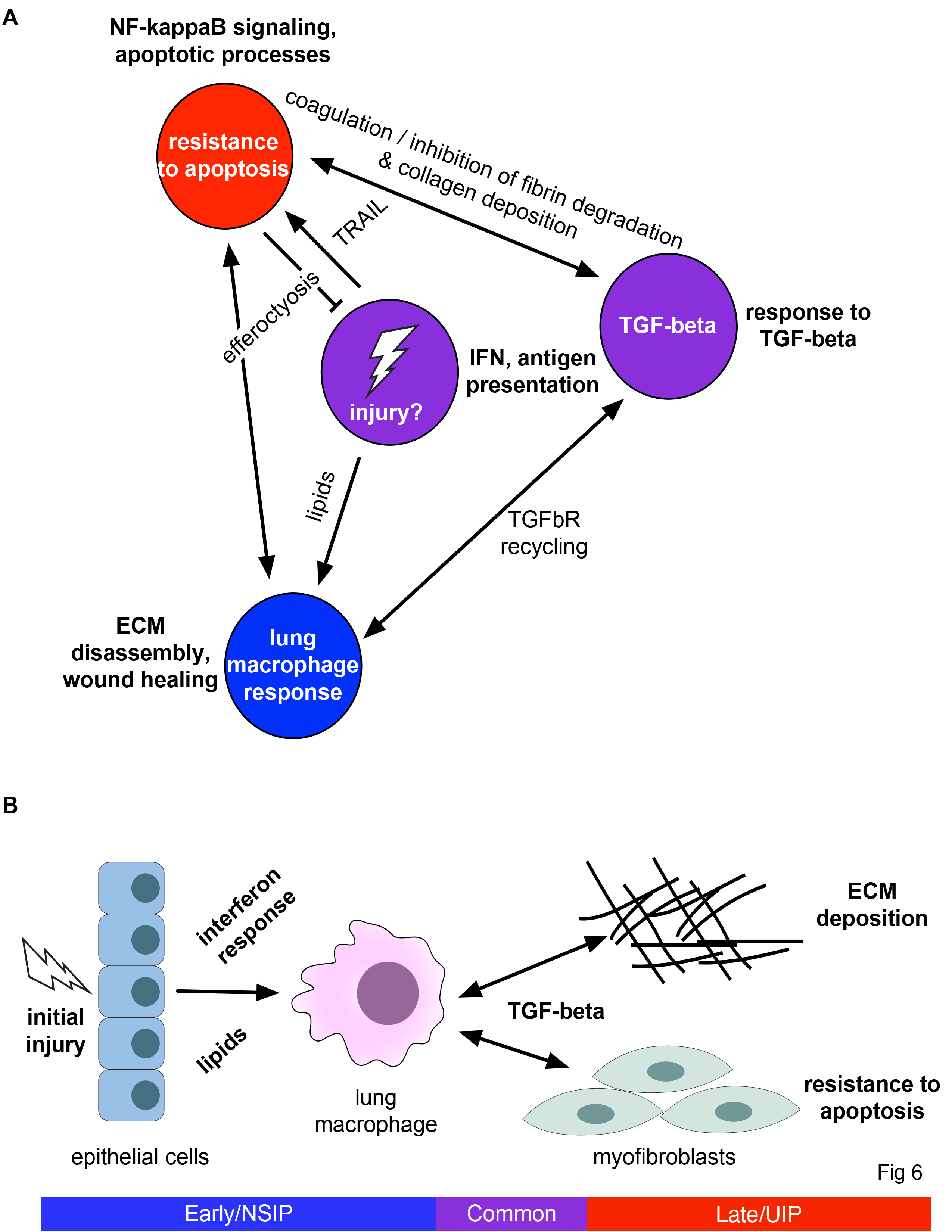
**Overview of SSc-PF disease processes**. (A) Network-centric overview (B) Cell type-centric overview.

Christmann and coworkers identified an increase in IFN-and TGF-β-regulated genes in biopsies from early SSc-PF (21). It was also noted that there was more CCL18 at the protein-level and a higher level of *CD163* transcript in SSc-ILD lungs, suggestive of the presence of alternatively activated M∅s (21). However, it was unclear which cell types were responsible for the IFN signature or if there was evidence of distinct subpopulations of M∅s. We found that gene signatures that are upregulated in alternatively-activated human M∅s and M∅s treated with free fatty acids are enriched in early SSc-PF patients and that there is no evidence for enrichment of a pro-inflammatory, IFN-stimulated M∅ signature (fig 5) (12).

The LR-M∅ signature identified in our differential network analysis consisted of genes with increased expression in early SSc-PF that participate in lipid and cholesterol trafficking (Figs 4D, S6, differential lung network). The expression of these genes is correlated with “canonical” M∅ genes identified in the primary publication (21) (fig 5). We find elevated gene expression programs associated with M∅ alternative activation (specifically metabolic “reprogramming”) and lipid exposure in this dataset (fig 5). In the bleomycin injury mouse model of pulmonary fibrosis, lipid-laden M∅s, or foam cells, have been observed to upregulate markers associated with alternative M∅ activation and to secrete TGF-β (59). Oxidized phospholipid treatment also causes alternative activation and TGF-β secretion in human M∅s (40). Consistent with this report, recent work demonstrates that foam cell formation *in vivo* favors the development of a pro-fibrotic M∅ activation profile (52,53). These studies, along with our results, suggest that lipid exposure or uptake in M∅s may be important.

TGF-β signaling is a hallmark of fibrotic disease, and was noted in the initial analysis of both lung datasets (20,21). Similarly, we find genes from both datasets in the response to TGF-β module of the lung network. However, we also find evidence that the type I IFN signature is present in the Bostwick dataset(fig 3). The functional module most strongly associated with late stage disease/UIP is the innate immune, NF-κB, and apoptotic processes module. This module is connected to the TGF-β module through components of the fibrinolysis pathway such as PAI-1 *(SERPINE1)* (fig 3). PAI-1 is upregulated in late stage SSc-PF and is known to be important in pulmonary fibrosis (54-56). One mechanism by which fibrinolysis may contribute to the resolution of fibrosis is through the induction of fibroblast apoptosis (57). Both TGF-β1 and PAI-1 have been shown to inhibit lung fibroblast apoptosis (57).

We found evidence for a shift in the balance of apoptosis in the Bostwick dataset, perhaps in myofibroblasts (58), in our network analyses (fig 6). Long-lived myofibroblasts are thought to continually deposit collagen and contribute to persistent fibrosis (59). This apoptotic-resistance phenotype is related to the stiffness of the matrix (60), suggesting that a shift in apoptotic processes may occur once the deposition of excess collagen begins. Moreover, impaired phagocytosis of apoptotic cells, or efferocytosis, has been observed in the alveolar M∅s of IPF patients (61). We find genes involved in efferocytosis, specifically in receptors *(CD44)* and endocytic machinery associated with this process, in the lung network (Figs 3, 6) (62). If the shift in apoptosis and efferocytosis occurs, we speculate that the fibrotic and inflammatory processes in our network will also be altered. Efferocytosis by alveolar M∅s plays a key role in the resolution of inflammation in the lung through the subsequent release of TGF-β (63). We hypothesize that following initial injury, TGF-β signaling, antifibrinolytic factors, and the disruption of apoptosis and efferocytosis may contribute to progressive fibrosis in SSc-PF (fig 6).

### Conclusions

In this study, we have utilized data from multiple tissues to examine the systemic nature of SSc. Our integrative analysis allowed us to leverage well-studied tissues to inform us about SSc manifestations that are under-studied molecularly. This study rigorously tests the notion that patients with severe disease have shared immunological and fibrotic alterations. The common immune-fibrotic axis shows evidence for alternatively activated M∅s in multiple SSc tissues. However, there are subtle differences in the M∅ gene expression programs detected in skin and lung. Different microenvironments likely provide distinct stimuli to infiltrating M∅s that determine the pro-fibrotic character of these cells. The plasticity of this lineage is likely central to the divergence of fibrotic processes in multiple SSc-affected tissues and is a central component of an immune-fibrotic axis driving disease.

## Methods

### Patients and datasets

Eight out of 10 datasets included in this study were previously published (see Table 1) and descriptions of the patient populations and criteria for inclusion can be found in those publications. We used the patient disease label (e.g., PAH) as published in the original work for all of these sets. In Table S1, we summarize the patient information to which we had access on a per array basis as that is what is required for comparison to the expression data. Below, we note some important characteristics (for the purposes of this work) of the included patient populations. As noted in the Results section, the two lung datasets contained patients with different histological patterns of lung disease. Some patients included in the PBMC dataset, including those with PAH, also had interstitial lung disease, though exclusion of these patients does not significantly change the interpretation as put forth in (18). As illustrated in S1 Table, two datasets (ESO, LSSc) did not contain healthy control samples and three datasets (UCL, LSSc, and PBMC) were comprised entirely of lcSSc patients.

### Ethics statement on previously unpublished datasets

The LSSc and UCL studies are previously unpublished. The samples from the LSSc dataset were obtained at Boston University Medical Center (BUMC)/Boston Medical Center (BMC); the BUMC/BMC Institution Review Board approved this study. The samples from the UCL dataset were obtained at University College of London; the London-Hampstead NRES Committee approved this study. The Dartmouth College CPHS approved this work. All subjects gave informed consent. All research conformed to the principles expressed in the Declaration of Helinski.

### Microarray dataset processing

This work contains 10 datasets on multiple microarray platforms. Agilent datasets (Pendergrass, PBMC, Milano, Hinchcliff, ESO, UCL, LSSc) used either Agilent Whole Human Genome (4×44K) Microarrays (G4112F)(Pendergrass, PBMC, Milano, Hinchcliff, ESO, UCL) or 8×60K (LSSc). Data were Log_2_–transformed and lowess normalized and filtered for probes with intensity 2-fold over local background in Cy3 or Cy5 channels. Data were multiplied by −1 to convert to Log_2_(Cy3/Cy5) ratios. Probes with >20% missing data were excluded. The Illumina dataset (Bostwick, HumanRef-8 v3.0 BeadChips) was processed using variance-stabilizing transformation and robust spline normalization using the lumi R package. Dr. Christmann provided the raw data in the form of . CEL files. Dr. Feghali-Bostwick provided Illumina BeadSummary files. Affymetrix datasets (Risbano, HGU133plus2; Christmann, HGU133A_2) were processed using the RMA method as implemented in the affy R package. Batch bias was detected in the ESO dataset. To adjust these data, missing values were imputed via *k*-nearest neighbor algorithm using a GenePattern (64) module with default parameters and the data were adjusted using ComBat (65) run as a GenePattern module to eliminate the batch effect.

To compare datasets in our downstream analysis, duplicate genes must not be present in the dataset and must be summarized in some way. First, we annotated each probe with its Entrez gene ID. Agilent 4×44K arrays were annotated using the hgug4112a.db Bioconductor package. LSSc was annotated using UNC Microarray Database with annotations from the manufacturer. Probes annotated to lincRNAs (A19) were removed from the analysis. The Illumina dataset was annotated by converting the gene symbols (provided as part of the BeadSummary file) to Entrez IDs using the org.Hs.eg.db package. The Risbano PBMC dataset was annotated using the hgu133plus2.db package. The Christmann dataset was annotated using an annotation file from the manufacturer. NAs and probes that mapped to multiple Entrez IDs were removed in all cases. Probes that mapped to the same Entrez ID were collapsed to the gene mean using the aggregate function in R, followed by gene median centering.

### Clustering of microarray data and statistical tests for phenotype association

The collapsed datasets were used to find coherent coexpression modules. We used Weighted Gene Co-expression Network Analysis (WGCNA), a strong clustering method, which allows us to automatically detect the number of coexpression modules and remove outliers (66). Each dataset was clustered using the blockwiseModules function in WGCNA R package using the signed network option and power = 12; all other parameters were set to default. The number of arrays and resulting co-expression modules are summarized in Table 2. Using the WGCNA coexpression modules also reduces the dimensionality of the dataset, as it allows us to test for genes′ association with or differential expression in, a particular pathophenotype of interest on the order of tens, rather than thousands using the module eigengene. The module eigengene is the first principal component, and represents the expression of all genes in a module and an idealized hub of the coexpression module. We used the moduleEigengenes function in the WGCNA R package to extract the eigengenes. A module was considered to be pathophenotype-associated if the module eigengene was significantly differentially expressed in or significantly correlated with a pathophenotype of interest. Only 2-class categorical variables were considered using a Mann-Whitney U test (i.e., all pulmonary fibrosis and pulmonary arterial hypertension patients were grouped together regardless of underlying etiology). We used Spearman correlation for continuous values. P-values were Bonferroni-corrected on a per phenotype basis. See S1 File for complete output. In the main text, we discuss categorical pathophenotypes, as these were enriched at the consensus cluster level. We do find instances coexpression modules that are associated with continuous pathophenotypes, such as pulmonary function test measurements, but these were not apparent at the consensus cluster level of abstraction.

### Module overlap network construction and community detection

The 10-partite ‘module overlap network’ was constructed as in Mahoney et al. (23), where it was called the ‘information graph’ due to its relationship to information theory. We describe the method here in brief and refer to (11) for motivating details. The modules from different datasets have no a priori relationship to each other. The module overlap network encodes the pairs of modules that significantly overlap. Specifically, for each pair of modules (C_i_ and C_j_) we compute an overlap score 
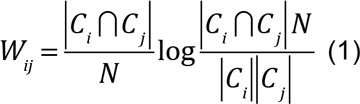
where N is the total number of genes shared between the two datasets. The overlap scores can be positive, negative, or zero, indicating that the modules overlap more, less, or the same as expected at random, respectively. As shown in Mahoney, et al. (11), the overlap scores can be naturally thresholded using information theory to yield a sparse network of significant overlaps. This is the module overlap network.

The module overlap network is highly structured. For example, a module representing an inflammatory process in skin often significantly overlaps inflammatory modules in other tissues. Thus, the structure of the module overlap network corresponds to the biological processes that are common to multiple datasets. We can identify these processes by clustering the module overlap network itself. To detect clusters in the module overlap network, we used two methods of community detection in the iGraph R package (67). First, we used fast-greedy modularity optimization (68), which yielded large, diffuse communities. We call these ‘top-level’ communities. To find smaller, more densely connected sub-communities, we used spin-glass community detection (igraph R package implementation, gamma.minus = 0.125, all other parameters were set to default) (67,69). We call these ‘bottom-level’ communities. The community/sub-community structure of the module overlap network demonstrates that there is a hierarchy of biological processes that are common across datasets, where large communities contain smaller ones (Fig. 2). To display this hierarchical community structure, we first sorted by top-level community label, and then within each community we sorted by bottom-level label. The adjacency matrix of the module overlap network and its node attributes (including dataset of origin and community labels) are supplied in S2 File.

We also tested each top-level community in the module overlap network for enrichment of pathophenotype-associated modules for each phenotype of interest using a Fisher’s exact test followed by Bonferroni correction (Table 5). This test takes into account both modules that had increased and decreased in pathophenotypes under study.

**Table 5.** **Bonferroni-corrected p-values, Fisher’s exact test pathophenotype-associated module in top-level communities in the module overlap graph**.

| Top-level community | 'In SSc' p-value | 'In Inflammatory' p-value | 'In Proliferative' p-value | 'In PAH' p-value | 'In PF' p-value |
| --- | --- | --- | --- | --- | --- |
| 1 | 1 | 0.02 | 1 | 1 | 1 |
| 2 | 0.71 | 0.07 | 1 | 1 | 1 |
| 3 | 0.09 | 0.27 | 1 | 0.77 | 0.29 |
| 4 | 8.56E-07 | 6.30E-12 | 1 | 0.30 | 1 |
| 5 | 1 | 1 | 0.03 | 1 | 1 |
| 6 | 1 | 1 | 1 | 1 | 1 |
| 7 | 1 | 0.64 | 1 | 0.03 | 1 |
| 8 | 1 | 1 | 1 | 1 | 1 |

### Functional and pathphenotype annotation of the module overlap network

The module overlap network contains rich information about the biological processes that are active in each tissue under study. We functionally annotated the module overlap network by finding pathways that strongly correlate to each community. Because an edge in the module overlap network corresponds to a significant overlap between coexpression modules from different datasets, we can think of an edge ‘encoding’ that overlap as a gene set. For each pair of coexpression modules *C_i_* and *C_j_,* we define an ‘edge gene set’, *E_ij_,* as the overlap between the in two datasets

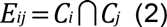

To annotate this edge gene set with biological pathways, we computed the Jaccard similarity of an edge gene set *E* and a pathway *P*

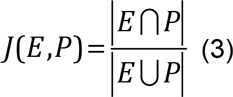

We used biological pathways from the Kyoto Encyclopedia of Genes and Genomes (70), BioCarta, and Reactome (71) obtained from Molecular Signatures Database from the Broad Institute (software.broadinstitute.org/gsea/msigdb). The Jaccard similarity between the edge and pathway will be equal to one, if all of the genes shared between two modules are exactly the same set of genes annotated to the pathway, or zero if no genes are shared between the two sets. To functionally annotate a community in the information graph, we compared the Jaccard similarities of the edges within the community to edges outside of the community using a Mann-Whitney U test (with Bonferroni adjustment). The full results of this analysis are included as S3 File.

### Tissue consensus gene sets

To understand how the immune and fibrotic responses in these phenotypes are functionally related, we found the consensus genes in the combined 4A and 4B clusters. Tissue consensus gene sets were derived by considering all modules within 4A and 4B, finding their unions within their dataset, and then computing their intersection across datasets from the same tissue of origin. For example, the lung consensus gene set (CC_lung_) was derived by computing the union of the Christmann (denoted *c*) and Bostwick (denoted *b*) modules in 4AB separately, and then computing the intersection across these two datasets:

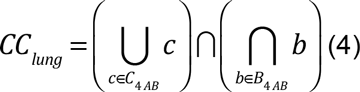

As each tissue was considered separately (limited skin and diffuse skin were considered separately), 5 tissue consensus gene sets were generated; the union of these tissue consensus datasets was used to query the functional genomic networks and is referred to as the ‘immune-fibrotic axis consensus’ gene set or genes throughout the text. For all genes in modules in clusters 4A and 4B, we calculated the Pearson correlation to their respective module eigengene (kME). We compared the kME of consensus genes to that of non-consensus genes using a Mann-Whitney U test. S3 Table contains the tissue consensus genes from 4AB or the ‘IMMUNE-FIBROTIC AXIS consensus genes.’

### Querying GIANT functional networks, single tissue network analysis, and network visualization

The GIANT functional genomic networks were obtained as binary (.dab) files and processed using the Sleipnir library for computational functional genomics (72). We queried all networks (lung, skin, ‘all tissue’) using the immune-fibrotic axis consensus gene sets (as Entrez IDs) and pruned all low probability (< 0.5) edges. All networks are available for download from the GIANT webserver (giant.princeton.edu) (1). For each single tissue analysis (consensus lung and consensus skin networks), we considered only the largest connected component of each network and performed spin-glass community detection as implemented in the igraph R package (67) to obtain the functional modules. We annotated functional modules using g:Profiler (73) using all genes in a module as a query. All networks in this work were visualized using Gephi (74). The network layout was determined by community membership, the strength of connections between communities, and finally the interactions between individual genes.

### Differential network analysis

The tissue-specific networks from GIANT allow for the analysis of the differing functional connectivity between genes in different microenvironments. In order to understand the specific immune-fibrotic connectivity in lung relative to skin, we performed a differential network analysis (fig 4). To compare networks we retained only nodes common to consensus skin network and consensus lung largest connected components (see above). We define the ‘differential lung network’ as the network with adjacency matrix: 
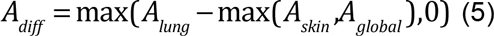
where A_lung_, A_skin_, and A_global_ are the lung, skin, and global (all tissues) adjacency matrices from GIANT. The differential lung network is thus the lung network minus the maximum edge weight from the skin and lung networks, where all edges that are stronger in skin or the global network are set to zero. Thus, the differential lung network contains only highly lung-specific interactions. Functional modules in the lung differential network were found using spin-glass community detection (see above) within the largest connected component of the network.

### Differential expression and M∅ gene set analysis

To identify genes that were differentially expressed in SSc-PF, SSc-PF samples were compared to normal controls in both datasets using SAM (23) (1000 permutations, implemented in samr R package). Genes with an FDR < 5% were considered further. The M∅ gene sets used in this study are WGCNA modules taken from a study of human M∅ transcriptomes (12). The z-score of each genes’ expression (Eqn. 6) was computed in the collapsed Christmann and Hinchcliff datasets (as described in ‘Microarray dataset processing’ section of Methods). The z-score *z* of gene *g* in the ith array/sample is computed as: 
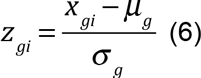
where *X_gi_* is the gene expression value in array/sample *i*, is the *μ_g_* gene mean, and *σ_g_* is the gene standard deviation. The average z-score of genes in a set (module from Xue, et al. (12)). computed for an array/sample to summarize gene set expression. Mann-Whitney U tests were used to compare average z-scores between groups (fig 5).

## Acknowledgements

JNT would like to thank members of the Whitfield Laboratory for thoughtful discussion.

## Funding

This work has been supported by grants from: the Scleroderma Research Foundation (SRF www.srfcure.org) to MLW and MH; grants from the Scleroderma Foundation to PAP and MLW; the Dr. Ralph and Marian Falk Medical Research Trust Catalyst Award to MLW; Dartmouth SYNERGY award to MLW; the Gordon and Betty Moore Foundation (GBMF4552) to CSG; National Institutes of Health (NIH; www.nih.gov) grants 2R01AR051089 to RL, P50 AR060780 and P30AR061271 to RL and MLW, K23 AR059763 to MH. JNT received support from NIH-NIGMS T32GM∅08704 and the John H. Copenhaver, Jr. and William H. Thomas, MD 1952 Junior Fellowship from Dartmouth Graduate Studies. CPD received support from the EULAR ODP programme. The funders had no role in study design, data collection and analysis, decision to publish, or preparation of the manuscript.

## Author Contribution

JNT, JMM, and MLW conceived of the study. JNT, CSG, VM, JMM, and MLW designed data analyses, performed analyses, and interpreted the results. TAW performed the microarray experiments. RBC, HWF, RAL, and CPD designed study cohorts included in this work and contributed samples and/or data. MEH provided clinical expertise and interpreted the results. PAP provided macrophage biology expertise and interpreted the results. JNT, PAP, JMM, and MLW wrote the paper. All authors read, revised, and approved the manuscript.

## Competing Interests

CPD has been a consultant to Roche, GlaxoSmithKline, Actelion, Inventiva, CSL Behring, Takeda, Merck-Serono, MedImmune and Biogen. MLW and MH have filed patents for gene expression biomarkers in systemic sclerosis. MLW is a scientific founder of Celdara Medical LLC. MLW has served as consultant to GlaxoSmithKline, Bristol Myers Squib, EMD Serono, Biogen and Quintiles. RL has received both grants and consulting fees from Genzyme/Sanofi, Shire, Regeneron, Biogen, BMS, Inception, Precision Dermatology, PRISM, UCB, Pfizer and Roche/Genentech; he received consulting fees from Lycera, Novartis, Celgene, Amira, Celdara, Celltex, Dart Therapeutics, Idera, Intermune, Medimmune, Promedior, Zwitter, Actelion, EMD Serono, Akros, Extera, Reneo, Scholar Rock, and HGS.

## Supporting Information Captions

**S1 Fig. Network view of consensus clusters 4A and 4B in the information graph**.

**S2 Fig. Density plot of correlation to respective module eigengene (kME)**. Tissue consensus genes have significantly higher kME and are therefore more ‘hub-like’ than non-consensus genes. Mann Whitney U, p < 2.2×10^−16^ reported by R

**S3 Fig. Fully labeled version of the consensus lung network (fig 3A**). This file is intended to be viewed digitally.

**S4 Fig. Fully labeled version of the differential adjacency matrix in Fig 4B**. This file is intended to be viewed digitally.

**S5 Fig. The differential lung network**. The highly lung-specific network (minus global network and skin network) contains functional modules.

**S1 Table. Table describing clinical characteristics of cohorts included in this study**.

**S2 Table. Consensus gene set sizes**.

**S3 Table. Immune-fibrotic axis consensus genes**.

**S4 Table. Mapping of Xue, et al. module numbers to our module names (Figure 5B**).

**S5 Table. P-values of all Xue, et al. modules tested**.

**S1 File. Tables - pathophenotype associations with WGCNA co-expression modules**.

**S2 File. Information graph adjacency matrix and module consensus cluster membership**.

**S3 File. Full output of edge-pathway (Jaccard) similarity Mann-Whitney U tests**.

**S4 File. Functional network edge lists and node attribute files (networks from Figures 3 and 4**).

The “common lung network” tab provides the module membership information for Fig 4B.

**S1 Text. Glossary of terms**.

**S2 Text. Additional results about pathophenotype-associated consensus clusters in the information graph**.

